# Translocon Remodeling Modulates Ribosomal Frameshifting and the Maturation of the Sindbis Virus Structural Polyprotein

**DOI:** 10.64898/2026.07.27.741003

**Authors:** Antonio Bonifasi, Gayani Apsara Ranasinghe, Ashish R. Jhangiani, Venkatesh P Thirumalaikumar, Brianna Corman, J. Paul Robinson, Suchetana Mukhopadhyay, Jonathan P. Schlebach

**Author notes:** Authors contributed equally (Antonio Bonifasi, Gayani Apsara Ranasinghe).

## Abstract

Like other enveloped RNA viruses, alphaviruses synthesize and assemble their envelope glycoproteins at the endoplasmic reticulum (ER) membrane. There, protein-conducting channels called translocons provide nascent proteins access to the membrane and allow them to fold into their correct shapes. We previously showed that a hydrophobic segment in the Sindbis virus structural polyprotein forms cotranslational interactions with the translocon that enhance −1 programmed ribosomal frameshifting (−1PRF), a recoding event that regulates polyprotein biogenesis. Recent discoveries concerning translocon remodeling suggest this segment, which corresponds to the second transmembrane domain of the E2 protein, could serve as a signal that recruits the multipass translocon (MPT). Here, we show that knocking out certain components of the MPT increases −1PRF efficiency. These differences in recoding coincide with changes in the membrane topology of the nascent polyprotein and in its downstream proteolytic processing in a manner that ultimately reduces viral fitness. Together, our results indicate that −1PRF and spike protein maturation in alphaviruses is tuned by the dynamic remodeling of the ER translocon. Such coupling could allow polyprotein biogenesis to adapt to different stages of viral replication and to distinct host or vector environments.

## Introduction

Alphaviruses are enveloped, positive-sense RNA viruses that are transmitted by arthropod vectors to small rodents, birds, and large mammals, including humans.^1–5^ The structural proteins of alphaviruses are translated from a subgenomic RNA as a single polyprotein consisting of the capsid, E3, E2, 6K, and E1 proteins, all encoded within one contiguous reading frame. The capsid protein is first translated and then autoproteolytically cleaved from the growing nascent chain.^6^ This cleavage event exposes a signal peptide at the N-terminus of the E3 protein, which prompts the delivery of the translating ribosome to the endoplasmic reticulum (ER) membrane. Elongation of the polyprotein then resumes at the ER, where the nascent chain matures through a series of post-translational cleavage, palmitoylation, and glycosylation reactions before it ultimately oligomerizes and traffics through the secretory pathway to the plasma membrane.^7–11^.

Although this structural polyprotein is the predominant translation product, alphaviruses also make use of a translational recoding mechanism known as -1 programmed ribosomal frameshifting (-1PRF) to modulate protein stoichiometry and expand their coding capacity. ^12,13^During translation, ∼15% of ribosomes transition into the -1 reading frame at a conserved slippery sequence (U_1_ UUU_4_ UUA_7_) within the 6K region, which results in the translation of the transframe (TF) protein in place of E1.^13–15^ This process is essential for maintaining the optimal stoichiometric ratio of the structural proteins that mediate viral assembly, and deviations in frameshifting can compromise infectivity and particle assembly.^16,17^

While -1PRF activity is typically attributed to certain *cis*-acting mRNA elements,^12,15,18^ we previously established that the processing of the nascent polypeptide chain allosterically stimulates -1PRF during the translation of the Sindbis virus (SINV) structural polyprotein.^19^ Like other alphaviruses, the SINV structural polyprotein features a marginally hydrophobic transmembrane domain within the E2 region (TMD2) that is inefficiently recognized by the translocon in a manner that promotes the formation of two competing topological isomers (Fig. 1A).^19^ In the major isoform, TMD2 bypasses the translocon and ultimately forms a loop connecting TMDs 1 & 3 (Fig. 1B), which is consistent with its orientation in the context of the assembled virion.^20^ The major isoform, which consists of the E3-E2-6K-E1 fusion protein (Fig. 1C), must adopt this orientation to ensure the E2 and E1 glycodomains are secreted into the ER lumen in order to facilitate the proper maturation and oligomerization of its spike proteins. However, the occasional translocon-mediated membrane integration of TMD2 creates a secondary topology and enhances -1PRF (Fig. 1D). This recoding event triggers the translation of the TF protein from the - 1-reading frame and the formation of the recoded polyprotein consisting of E3-E2-TF, respectively (Fig. 1E). These cotranslational interactions coincide with ribosomal stalling on the -1PRF slip-site, which is mediated by an mRNA stem-loop.^21^ Decoupling these processes greatly reduces -1PRF,^19^ which demonstrates the allosteric nature of this crosstalk.

**Figure 1.**
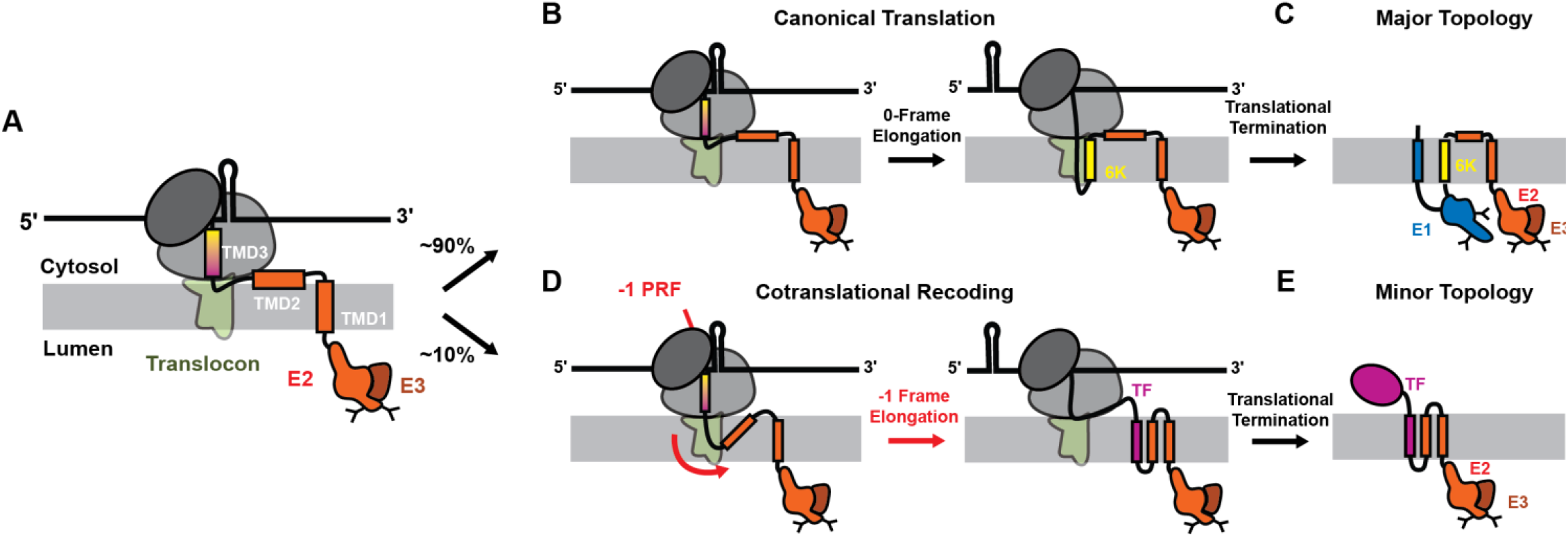
Biosynthesis of Sindbis Structural Polyprotein Isoforms. Cartoons depict the relationship between the cotranslational processing and the assembly competence of the major and minor isoforms of the SINV structural polyprotein (schematics are not to scale). A) As the nascent chain emerges at the ER membrane, TMD2 of E2 either bypasses the translocon and forms a cytosolic loop (major pathway), directing continued translation in the 0-frame, or is integrated into the lipid bilayer (minor pathway), which stimulates −1 programmed ribosomal frameshifting (−1PRF) and directs translation into the −1 frame. Under the conditions used here, the minor pathway accounts for approximately 10% of translation events in parental HEK293T cells (Fig. 3F).^19^,^21^ B) In the major pathway, TMD2 remains cytosolic as a loop connecting TMD1 and TMD3, and elongation proceeds through the 6K region in the 0-frame. C) The resulting major isoform is a nascent E3-E2-6K-E1 fusion protein that adopts an orientation in which the E1 and E2 glycodomains reside in the ER lumen, positioning the downstream cleavage sites for proteolytic maturation. D) In the minor pathway, membrane integration of TMD2 inverts the orientation of the downstream sequence and stimulates −1PRF, after which elongation continues in the −1 frame through the transframe (TF) open reading frame. E) The resulting minor isoform is a nascent E3-E2-TF fusion protein in which the E2 glycodomain resides in the ER lumen and TF is projected into the cytosol.

The link between nascent-chain topology and translational recoding appears to stem from mechanochemical allostery within the ribosome-translocon complex.^21^ Earlier biochemical measurements and coarse-grained simulations suggested that both the abundance of the minor topomer and the net −1PRF efficiency are coupled to specific interactions with the Sec61 translocon.^21^ However, this interpretation is cofounded by recent findings suggesting the same topological features within this nascent polyprotein may also stimulate the dynamic remodeling of the translocon.^22–25^ The E2 protein contains two TMDs that are linked by a short loop, which is a feature known to trigger the recruitment and assembly of the multipass translocon (MPT).^23^ In the MPT complex, Sec61 acts as a scaffold that associates with the PAT (CCDC47/Asterix), BOS (Nicalin/TMEM147/NOMO1), and GEL (TMCO1/OPT1) accessory complexes.^26^ Indeed, the MPT serves as the primary translocon that facilitates the cotranslational assembly of other multipass membrane proteins.^24,25^ Together, these observations suggest that translational recoding in the SINV structural polyprotein could potentially coincide with the dynamic remodeling of the translocon.

In the following, we assess the impact of the MPT on viral protein biogenesis. We first show that depletion of certain translocon subunits significantly increases -1PRF efficiency. We then show that disrupting the MPT enhances the formation of the minor topomer and reduces the maturation efficiency of the structural polyprotein. Finally, we demonstrate that these observed changes in recoding and topology result in the attenuation of viral fitness. Together, these findings reveal that translational recoding in the SINV structural polyprotein is dynamically modulated by the composition of translocons within the host cell.

## Results

### Characterization of Multipass Translocon Knockout Cells

To assess the role of the MPT in polyprotein biogenesis, we used CRISPR/Cas9 to generate a series of knockout cell lines lacking some of its core components. We targeted the *CCDC47, TMCO1*, or *NOMO1* genes, which encode core subunits of the PAT, GEL, and BOS accessory subunits, respectively. In addition to confirming genomic knockout (Fig. S1), western blots and proteomic characterization demonstrated that the target proteins and certain binding partners within the PAT, BOS, and GEL subunits are depleted in the knockout cells (Figs. 2 & S1). Similar co-depletion of partner subunits has been previously reported in knock down cells.^23^ To assess how the loss of these proteins remodels cellular translocons and other potential host protein interactors, we quantitatively compared the proteome of each knockout cell line. As expected, an analysis of the relative abundances of over 9,500 proteins in each cell line revealed that CCDC47, NOMO1, and TMCO1 are among the proteins that exhibit the large decreases in abundance in their respective knockout lines (Fig. 2A-C). Nevertheless, our proteomic analysis also identifies over 50 other proteins in each knockout line that exhibit at least a 2-fold variation in abundance relative to the parent (Table S1), including some components of secondary translocons such as the ER membrane protein complex (EMC, Fig. 2D). Nevertheless, it should be noted that these secondary effects of MPT depletion did not appear to complicate previous investigations of MPT structure and function.^22,23^ Moreover, the core components of the Sec61 translocon and Oligosaccharyltransferase (OST) complex remain intact in these cells (Fig. 2D), indicating that the general secretory machinery persists upon depletion of the PAT, BOS, and GEL complexes.

**Figure 2.**
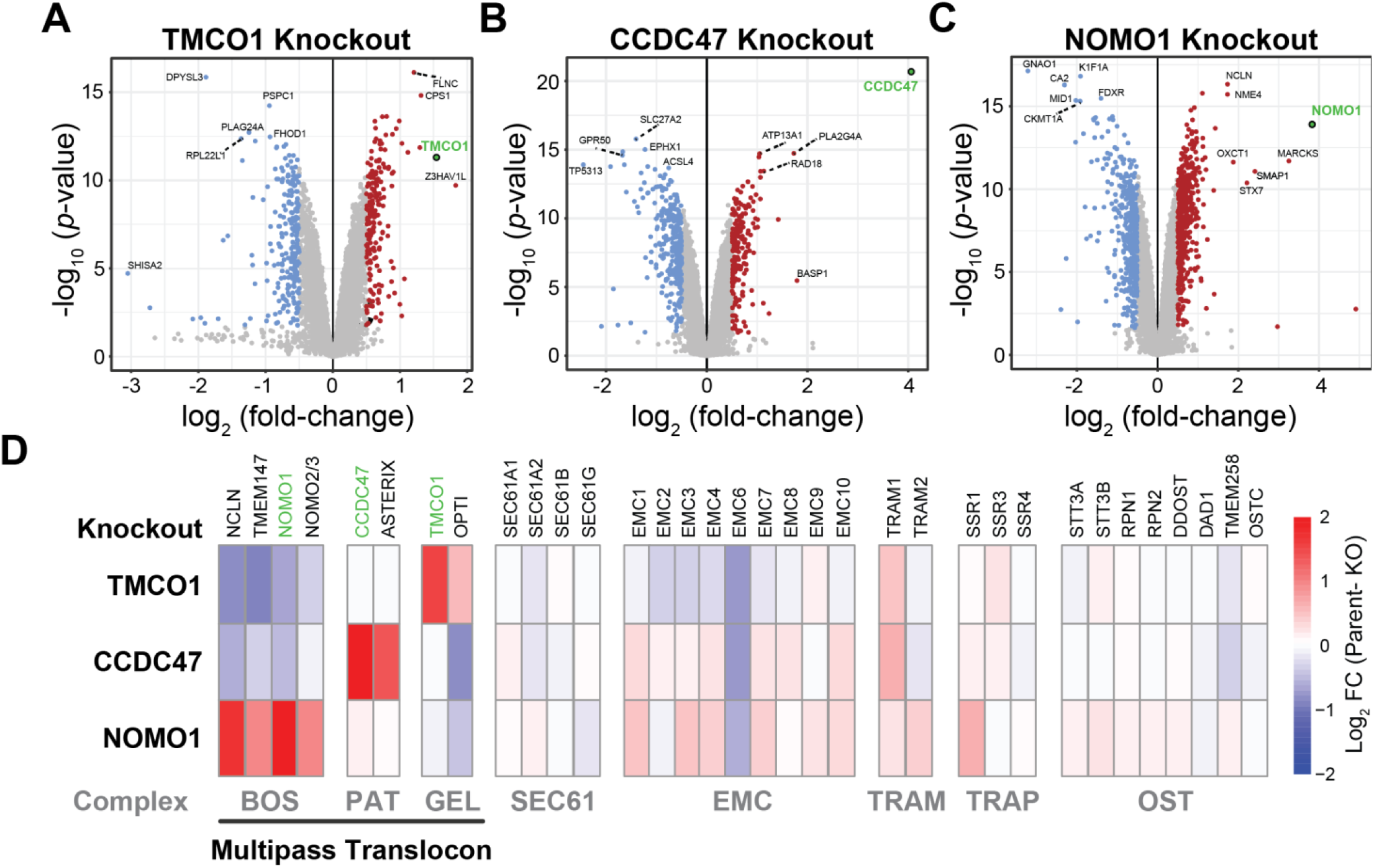
Proteomic Characterization of Multipass Translocon Knockout Cells. Whole-cell proteomics was used to quantitatively compare changes in the abundances of cellular proteins in the context of CRISPR knockout cell lines lacking components of the multipass translocon. A series of volcano plots depicts the log_2_ fold-change in abundance (parent – knockout) for each identified protein relative to the corresponding -log_10_ of the corresponding *p*-value in the context of A) TMCO1, B) CCDC47, and C) NOMO1 knockout cells relative to the parental HEK293T cell line. Statistical significance was assessed using a student’s t-test. D) A heatmap depicts the log_2_ fold-change values associated with subunits of various translocon complexes in the context of each knockout cell line. Red indicates a reduced abundance of corresponding protein in the knockout cell line whereas blue indicates an increase in abundance. Values represent the average across four biological replicates.

### Modulation of -1PRF by the Multipass Translocon

To determine whether the recruitment of the multipass translocon modulates -1PRF, we used a previously established fluorogenic genetic reporter^19^ to compare -1PRF efficiency across this panel of knockout cell lines. This reporter features a 1656 bp fragment of the sub-genomic viral RNA beginning with the region encoding the E3 signal peptide and continuing through the structural RNA elements of the -1PRF site (Fig. 3 A-B). Importantly, this design ensures the E3 signal peptide emerges first from the ribosome, a key consideration to preserve its SRP-mediated targeting to the ER membrane. Downstream in the reporter, the viral -1PRF site is followed by a -1 GFP cassette that is selectively translated in response to a -1 ribosomal frameshift and is released from the polyprotein via a self-cleaving 2A linker (Fig. 3 A-B). Finally, the frameshift reporter is followed by an internal ribosome entry site (IRES) that promotes the constitutive translation of a fluorescent mKate expression marker. We transiently expressed this reporter in each cell line, then utilized flow cytometry to measure single-cell fluorescence profiles. To control for variations in reporter expression, we used cellular mKate intensities to restrict our analysis to a subset of cells within a discrete reporter expression level (Fig. 3C). Notably, cells with this equivalent expression level exhibit GFP levels above cells expressing various negative control constructs that lack a slip-site or that contain stop codons in either reading frame (Fig. S2), which confirms the GFP signal is generated by a -1 ribosomal frameshift. GFP levels within these populations exhibit significant variations, suggesting -1PRF efficiency is sensitive to the loss of the multipass translocon (Fig. 3D). To quantify variations in -1PRF efficiency, we first normalized the -1PRF GFP signal by cellular mKate intensities (Fig. 3E), then calculated the absolute frameshifting efficiency using a control in which GFP is translated from the 0-frame-an approximation for the maximum GFP signal (Fig. S2). Overall, -1PRF efficiency trends higher in each knockout cell line relative to the parent, with a statistically significant increase occurring in the CCDC47 cell line (Fig. 3F). The -1PRF increases from 10 ± 2 % in the parental HEK293T cell line to 14 ± 2% in CCDC47 knockout cells (*p* = 0.016, Fig. 3F). These results demonstrate that translocon remodeling significantly alters translational recoding in the SINV structural polyprotein. Our findings suggest the recruitment of certain components of the MPT, especially the PAT complex, typically suppresses ribosomal frameshifting during spike protein biogenesis.

**Figure 3.**
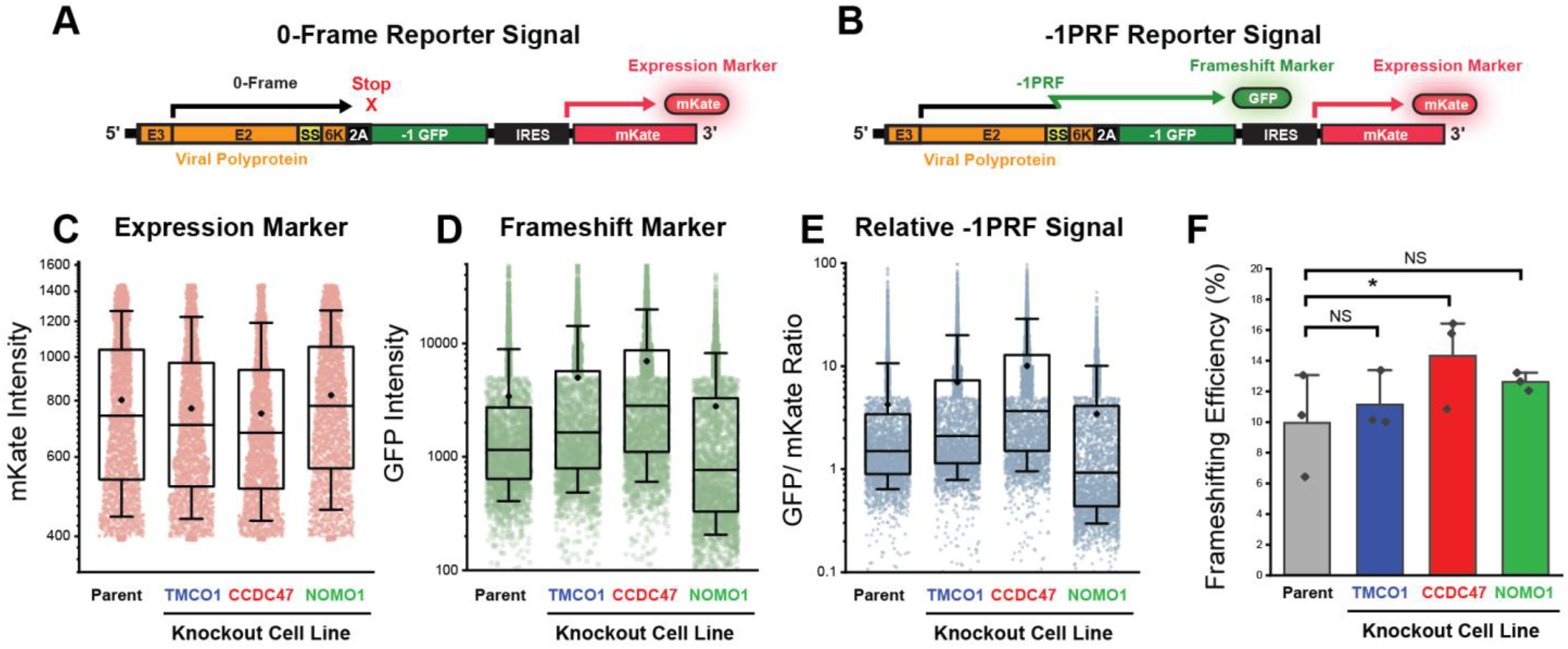
Impact of the Multipass Translocon on -1 Programmed Ribosomal Frameshifting in the SINV structural polyprotein. A genetic -1PRF reporter that generates both a constitutive mKate expression marker and a GFP in response to -1PRF in the SINV structural polyprotein was transiently expressed in a series of HEK293T knockout cell lines lacking TMCO1, CCDC47, or NOMO1 prior to analysis of cellular fluorescence proteins by flow cytometry. Cartoon diagrams of the reporter show how A) translation in the 0-frame generates mKate while B) -1PRF generates both GFP and mKate. Box and whisker plots depict the range of single-cell C) mKate intensities, D) GFP intensities, and E) GFP/ mKate ratios among cells within a discrete expression level across the knockout cell lines for a representative experimental replicate. Upper and lower bounds of the boxes reflect the 75^th^ and 25^th^ percentile values, respectively. The upper and lower whiskers reflect the positions of the 90^th^ and 10^th^ percentile values, respectively, and the midlines represent the median. F) A bar graph depicts average -1PRF efficiencies in each cell line and the whiskers reflect the standard deviation across four biological replicates. A student’s t-test was used to determine whether the difference in the means is statistically significant (* *p* < 0.05).

### Modulation of Polyprotein Topology by Multipass Translocon Subunits

We previously showed that variations in -1PRF efficiency can arise from differences in TMD2 translocation.^19^ However, previous experiments were carried out in HEK293T cells containing the full complement of translocon subunits. To assess whether changes in recoding coincide with differences in topology, we used a series of previously described topology reporters to detect changes in the orientation of the nascent polyprotein MPT knockout cells.^19^ These reporters feature a glycosylatable GFP (gGFP) domain, which fluoresces in the cytosol but not in the ER lumen, inserted at various positions downstream of each TMD (Fig 4 A-B). Notably, the membrane integration of TMD2 in the minor topomer inverts the orientation of TMD3 in a manner that should increase the gGFP signal of the TMD3 reporter and the TMD3+ reporter, which includes an additional soluble residues downstream of the TMD (Fig. 4 A-B).

**Figure 4.**
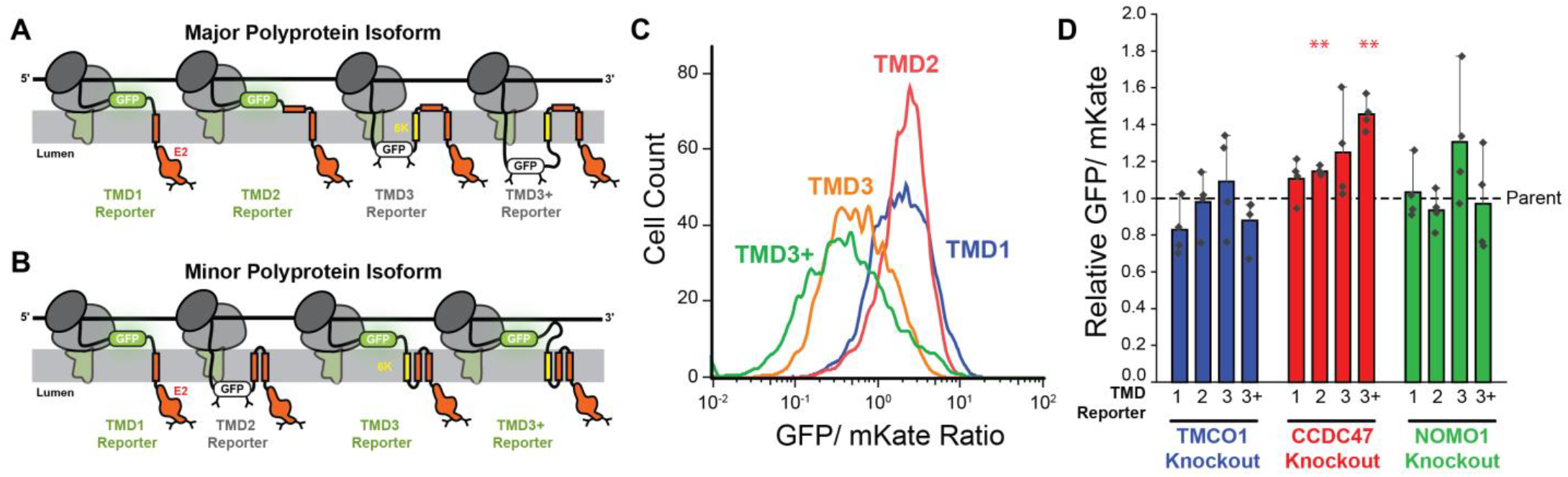
Impact of the Multipass Translocon on Polyprotein Topology. A series of genetic reporters were used to probe the topological orientation of the nascent SINV structural polyproteins in knockout cell lines. Cartoons depict the compartmentalization of the glycosylatable GFP sensor in the context of A) the major topological isoform and B) the minor topological isoform for each successive reporter construct. C) A histogram depicts the distribution of cellular GFP: mKate ratios among parental HEK293T cells transiently expressing the TMD1 (blue), TMD2 (red), TMD3 (orange), and TMD3+ (green) topology sensor constructs for a representative replicate. D) A bar graph displays the relative GFP/ mKate ratios among TMCO1 (blue), CCDC47 (red), and NOMO1 (green) knockout cells transiently expressing each reporter construct. Values were normalized according to the corresponding ratio in the parental HEK293T cell line. Values represent the average across four biological replicates, and whiskers reflect the standard deviation. Statistical significance was determined using a student’s t-test (* *p* < 0.05, ** *p* <0.01).

We expressed this collection of reporters in each knockout line, then quantitatively compared trends in gGFP fluorescence by flow cytometry (Fig. 4C). As before, we selected cells based on a discrete intensity interval associated with a constitutive IRES-mKate expression reporter in order to control for variations in transfection, then used its signal to normalize single-cell gGFP reporter intensities. Overall, the TMD1 and TMD2 reporters generate higher gGFP: mKate ratios relative to the TMD3 and TMD3+ reporters in each cell line (see Fig. 4C), which reflects the dominant contribution of the major isoform in which the C-terminus is translocated across the ER membrane by TMD3 (Fig. 4A). However, subtle variations in intensity were detectable in certain cases. Quantitative trends in cellular gGFP: mKate ratios within TMCO1 and NOMO1 knockout cell lines are statistically indistinguishable from those in parental cells (Fig. 4D). In contrast, cells expressing the TMD3+ reporter exhibit 46 ± 9% higher cellular gGFP: mKate ratios in CCDC47 knockout relative to parental HEK293T cells expressing this reporter (*p* = 0.003, Fig. 4D). A statistical difference was also observed for the TMD2 reporter in this cell line, though the magnitude of the difference is relatively small. As expected, the overall shifts in the nascent topological ensemble are subtle given the ∼10-20% relative abundance of the minor topomer.^19^ Nevertheless, the relative strength of the phenotype in CCDC47 knockout cells relative to NOMO1 and TMCO1 knockout cells is consistent with trends in the -1PRF reporter (Fig. 3). Together, these results suggest that the depletion of the PAT complex increases the abundance of the minor isoform and enhances -1PRF.

### Impact of the Multipass Translocon on Structural Polyprotein Processing

Proper proteolytic processing of the SINV structural polyprotein requires that its cleavage sites are localized within their correct cellular compartments. Thus, the observed variations in polyprotein biogenesis in knockout cells could potentially reduce the efficiency of polyprotein maturation by altering the exposure of proteolytic cleavage sites. To compare the polyprotein maturation efficiency across the panel of knockout cells, we infected these cells with SINV and used western blotting to compare the processing efficiency of the polyprotein within each cell line. Briefly, cells were infected at a multiplicity of infection (MOI) of 1.0 then harvested after 48 hours. Cell lysates were run on an SDS-PAGE and a western blot using an anti-E2/E1 primary antibody, which detects three polyprotein states, including higher-order structures along with the immature p110 precursor protein (E3-E2-6K-E1) and the mature E2/E1 proteins (Fig. 5A). Based on the relative abundance of the p110 compared to E2/E1 glycoproteins, we observe minimal difference in the production and/ or processing efficiency of the polyprotein in the TMCO1 knockout line. In contrast, we observe significantly lower protein levels and attenuated processing efficiencies in the CCDC47 knockout line relative to the parental HEK293T cells. NOMO1 knockout cells exhibited a mixed response, with lower cleavage efficiency relative to parental cells, though the differences in the means are not statistically significant (Fig. 5B). Together, these findings confirm that disrupting the MPT, in particular the PAT complex, compromises translational recoding, membrane integration, and polyprotein processing.

**Figure 5.**
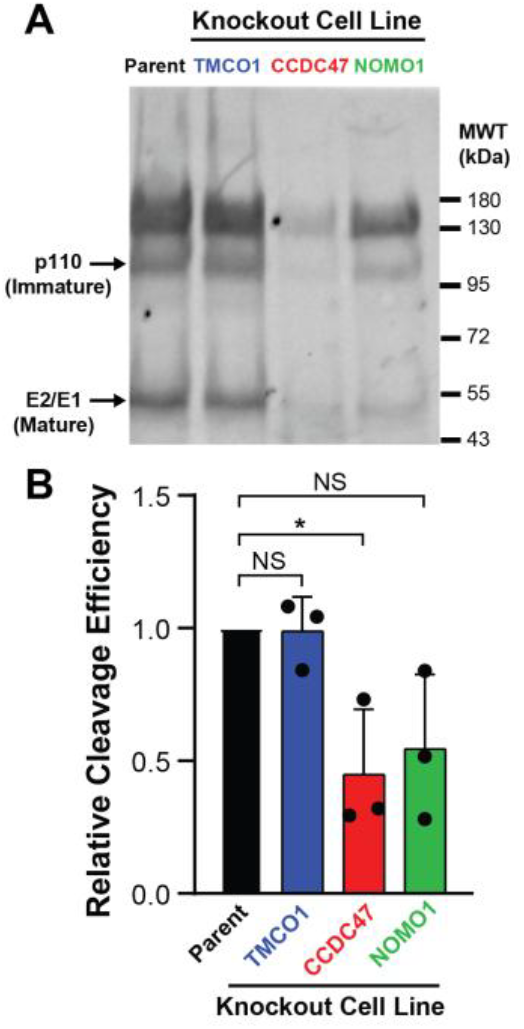
Impact of the Multipass Translocon on Structural Polyprotein Maturation. The processing efficiency of the SINV structural polyprotein is compared across the parental and knockout HEK293T cells. A) A representative western blot against the SINV E2/E1 proteins depicts the relative abundance of immature poly-proteins (p110, E3-E2-6K-E1) and mature E1 proteins within cell lysates collected after infection of parental and knockout cell lines with the SINV virus (MOI = 1). B) A bar graph depicts the relative polyprotein cleavage efficiency in each cell line (n = 3). Statistical significance was assessed using an unpaired, two-tailed t-test (\**p* < 0.05).

### Impact of Multipass Translocon Subunits on Viral Fitness

Polyprotein maturation efficiency ultimately dictates the relative abundance of mature viral proteins that are competent for assembly. Thus, decreases in processing efficiency could reduce viral fitness. To assess how differences in polyprotein maturation impact viral infection, we compared the SINV propagation kinetics across the panel of MPT knockout cells. We first used a modified version of the SINV TE12 strain containing a self-cleaving mCherry-FMV2A between the CP protein and the E3 signal peptide to track viral infection. Cells were infected at a multiplicity of infection (MOI) of 1.0 and imaged in real-time for phase and red fluorescence every 3-6 hours (Fig. 6A). In parental HEK293T cells, the onset of infection was observed at 14.3 hours (Fig. 6C), reaching 50% infection by 24 hours and peaking at 36 hours (Fig. 6 A & D). Though infection follows a similar time course in TMCO1 knockout cells, viral spread was delayed in both the CCDC47 and NOMO1 knockout cells, with the greatest attenuation occurring in the CCDC47 knockout (Fig. 6A). Indeed, the onset of infection was delayed to 25 hours in CCDC47 knockouts, with 50% infection occurring at 40 hours and peak infectivity requiring 61.5 hours. Similarly, the NOMO1 knockout cells exhibited delayed 50% infection mark at 32.5 hours and peaked at 54.5 hours (Figs. 6C-E). We corroborated these findings by measuring viral titers (Log_10_ pfu/ml) over the same time course. Consistent with fluorescence data, the viral titers demonstrated that the CCDC47 and NOMO1 knockout lines yielded substantial lag in viral titer relative to the parental and TMCO1 knockout cells (Fig. 6B). Nevertheless, viral titers converge after 40 hours (Fig. 6B), suggesting the viruses eventually accumulate a sufficient quantity of mature polyprotein to assemble, even without the MPT. Notably, several mutant SINV strains bearing mutations that disrupt the native frameshift elements also exhibit fitness defects in the MPT knockout lines (Fig. S3), suggesting the reduced fitness may arise from defects in spike protein assembly rather than variations in -1PRF. These results confirm that the attenuated polyprotein maturation in the PAT knockout cells reduces viral fitness. Collectively, our findings across reporter measurements and viral infection data cumulatively suggest a loss of MPTs disrupts polyprotein processing and translational recoding in a manner that reduces viral fitness.

**Figure 6.**
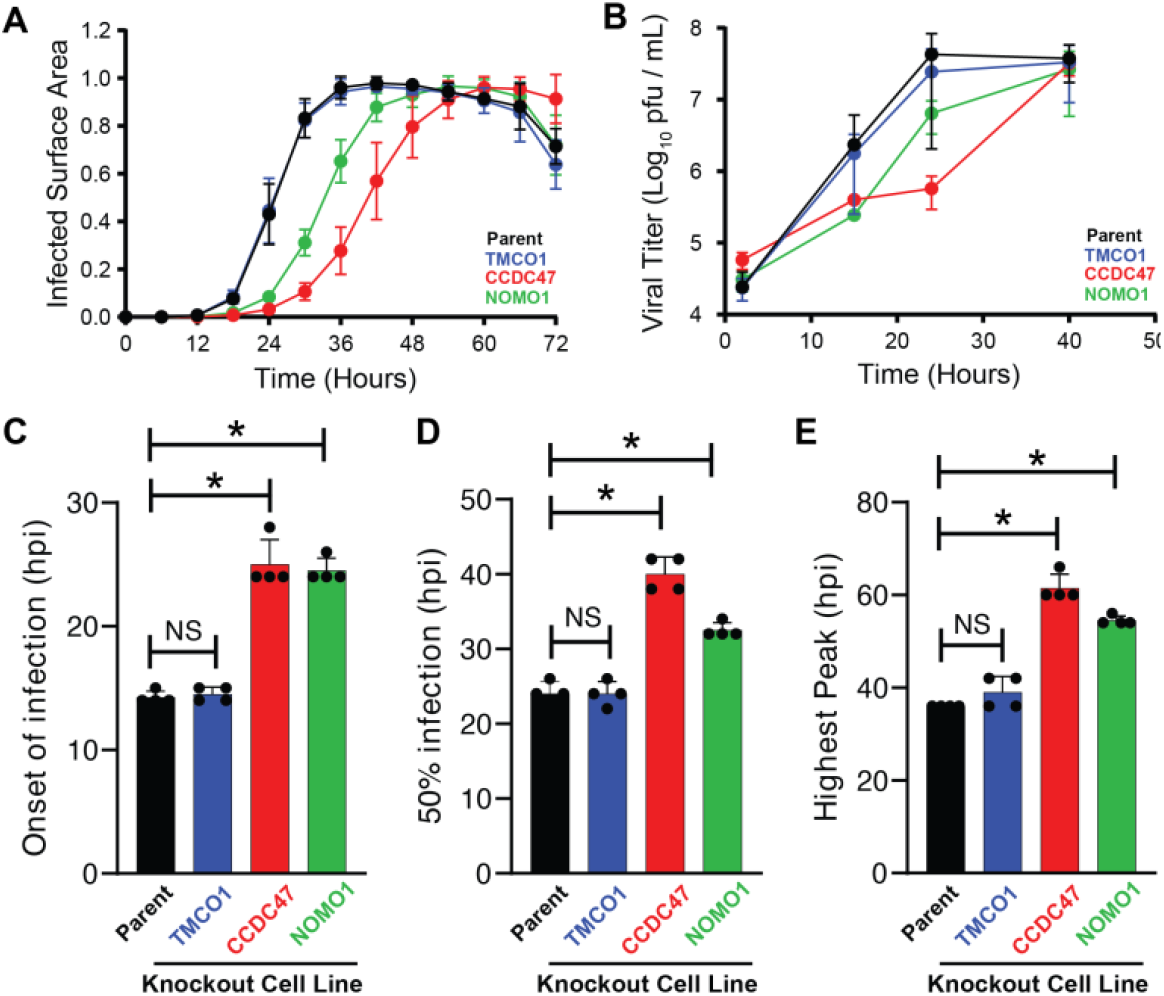
Impact of the Multipass Translocon on Viral Fitness. The growth kinetics and infectivity of viral particles are compared across parental and knockout cells. A) The spread of an mCherry-tagged SINV was tracked over time using an Incucyte live cell imager. The relative surface area of cells infected at an MOI of 1.0 is plotted over time for NOMO1 knockout (green), CCDC47 knockout (red), TMCO1 knockout (blue), and parental HEK293T cells (black). Data are normalized to the highest fluorescent value in each cell line. B) Viral infectivity was tracked over time by measuring the number of infective viruses within the growth media at various points after the initiation of infection. The concentration of plaque-forming units is plotted against the time after the initiation of infection at an MOI of 1.0. Points represent the average of three biological replicates and error bars reflect the standard deviation. Time to 50% infection C), onset of infection D), and time to peak titer E), calculated from A) for each cell line. Points represent biological replicates; bars show mean ± SD. Statistical significance determined by two-tailed t-test comparing knockouts to parental (*p < 0.05; NS, not significant; n = 3).

## Discussion

Though viral -1PRF has historically been attributed to the activity of *cis*-acting mRNA elements,^18^ we previously established that mechanical feedback generated by membrane translocation stimulates recoding in the SINV structural polyprotein. At the time of these discoveries, Sec61 was generally believed to independently establish the topology of multipass membrane proteins.^27^ However, recent discoveries have shown that it is instead dynamically remodeled throughout translation in order to accommodate the assembly of a diverse array of membrane protein folds.^22–24^ Our findings reveal that this remodeling coincides with a pivotal transition in the biogenesis of the SINV structural polyprotein, at which the ribosome commits to translational recoding and the production of a frameshifted virulence factor. We show that certain components of the MPT impact the topology and processing of the nascent polyprotein in manner that coincides with variations -1PRF efficiency. Recoding measurements in CCDC47 knockouts demonstrate that the loss of the PAT complex increases -1PRF efficiency (Fig. 3) and enhances the formation of the minor topological isomer (Fig. 4). This, in turn, compromises the maturation of the E2 and E1 spike glycoproteins (Fig. 5) and ultimately attenuates viral fitness (Fig. 6). Together, these results implicate certain components of the MPT in the allosteric regulation of the translational recoding that tunes the stoichiometry and processing of viral structural proteins.

There are several limitations of this study that warrant consideration. We first note that the limited sensitivity of our genetic reporters makes it challenging to resolve some of the finer aspects of polyprotein topology and recoding given its ∼10-20% efficiency (Fig. 3F). While we observe consistent trends in the effects of knockouts across divergent experiments (Fig. 3-6), the effects are generally only robust enough to consistently achieve statistical significance in CCDC47 knockout cells. Unfortunately, the low frequency and high complexity of these interactions also make it challenging to directly interrogate the biochemical and/ or biophysical aspects of this mechanism. Second, it should be noted that, while the allosteric feedback involved in recoding is a discrete, kinetically controlled event,^28–30^ our use of stable knockout cells can only capture the collective steady-state consequences that arise from bulk perturbations of these translocon complexes. Indeed, these knockout cells exhibit non-equivalent proteomic variations, including differences in the expression of the EMC insertase (Fig. 2, Table S1). These differences could create pleiotropic effects that confound our interpretations. Nevertheless, similar caveats should hold true for other cellular and/ or biochemical investigations that rely on rough microsomes from CRISPR knockout cells for *in vitro* translation.^23^ Finally, we note that HEK293T cells are deficient in type I interferon signaling. Thus, our viral experiments do not capture any immunomodulatory consequences of altered TF production that may shape viral fitness.^31^ Despite these caveats, our findings clearly establish a mechanistic connection between translocon remodeling and translational recoding in SINV.

The results described herein suggest that recruitment of certain translocon subunits alters nascent chain interactions in a manner that ultimately modulates translational recoding. Measurements in CCDC47 knockout cells show that the loss of the PAT complex enhances recoding efficiency and the corresponding production of the minor topomer where TMD2 is integrated into the membrane (Figs. 3-4). This implies that the recruitment of the MPT coincides with the shunting of TMD2 into the cytosol, the projection of the E1 and E2 glycodomains into the ER lumen, and the translation of E1 (Figs. 1 & 7). In conjunction with observations,^19,21^ our current findings suggest translocon composition potentially tunes the tension on the nascent chain in a manner that modulates recoding efficiency. Notably, a previous deep mutational scan identified a cluster of hydrophobic residues in the C-terminal portion of TMD2 that is essential for coupling between translocation and -1PRF.^21^ We previously speculated that this arose from specific interactions between the nascent polypeptide and the lateral gate of Sec61 in the context of the minor isoform. Our current findings potentially suggest these same residues may participate in the recruitment of PAT and/ or other MPT components in the context of the major isoform (Fig. 7). We note that the pronounced phenotypes in PAT knockout cells may suggest the translocons engaged at the point of frameshifting may lack BOS and/ or GEL. While current evidence suggests these three complexes act in tandem as an obligate oligomer,^22–25^ an independent role of PAT^32^ seems plausible given that -1PRF is a kinetically controlled event that may occur at a specific intermediate stage of MPT assembly.

**Figure 7.**
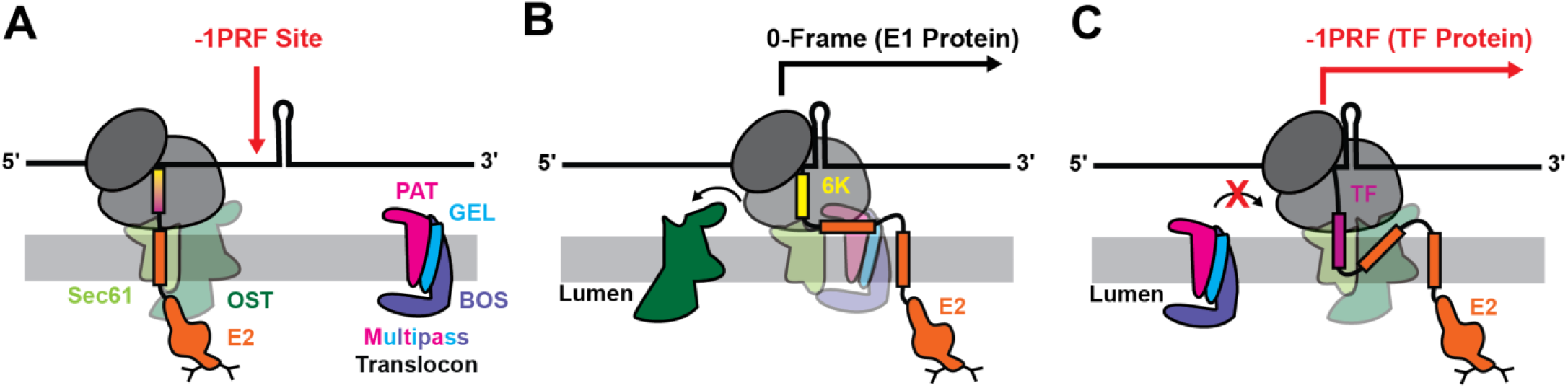
Translocon Remodeling Modulates Translational Recoding. A cartoon outlines how the remodeling of the translocon during SINV structural polyprotein biosynthesis influences translational recoding. A) The ribosome is associated with the oligosaccaryltransferase subunit (OST) as the ribosome approaches the -1PRF site while synthesizing the second TMD of the E2 protein. B) The emergence of the second TMD from the ribosomal exit tunnel triggers the assembly of the multipass translocon, which displaces the OST and suppresses -1PRF. Elongation then continues in the 0-frame, which results in the synthesis of the E1 spike protein. C) Under conditions in which the multipass translocon is not efficiently recruited during recoding, the translocation of the second TMD occurs while the ribosome is stalled on the slip-site, which triggers -1PRF. Elongation then continues forward in the -1 frame, which results in the synthesis of the TF protein.

Overall, our observations also serve to highlight translocon subunits as important host factors involved in viral assembly. Variations in the abundance of available translocons clearly modulates the accumulation and maturation of structural proteins (Figs. 5-6). Elevated −1PRF in the absence of the MPT enhances TF production at the expense of E1 and 6K, which in turn reduces the yield of assembly-competent spike proteins. Thus, the availability of multipass components could potentially constrain viral infectivity. Since the MPT assembles on-demand around multipass substrates^24^ and its subunits are likely present at sub-stoichiometric levels relative to Sec61,^25^ the pool of assembled complexes is necessarily finite. Such complexes may become scarce as infection progresses and the cell upregulates the translation of viral spike proteins. This raises the possibility that -1PRF and TF production may primarily occur during the late stages of infection, when spike proteins have already matured and assembled. The availability of certain translocons may also constrain the ability of certain alphaviruses to spread between certain hosts and vectors. While the core subunits of the MPT are conserved across eukaryotes, it is possible that the composition and activity of translocons may vary in a manner that differentially modulates recoding in mosquitos and other invertebrate hosts. Additional investigations are needed to evaluate how translocon remodeling constrains viral infection.

## Materials and Methods

### Cell Culture

HEK293T-derived cells were grown in Dulbecco’s modified eagle medium (Gibco, Carlsbad, CA) supplemented with 10% fetal bovine serum (Corning, Corning, NY) and penicillin (100 U/mL)/streptomycin(100 µg/mL) at 37°C in a humidified incubator containing 5% CO_2_ by volume.

### Reporter Construct Design and Preparation

The bicistronic -1PRF reporter construct used in this study was generated as previously described.^15^ Briefly, a fragment of the SINV sub-genomic RNA spanning from the E3 signal peptide through 100bp downstream of the ribosomal slip site was cloned into a vector upstream of a GFP cassette encoded on the –1-frame. An IRES-mKate cassette was added downstream of the test construct to serve as an expression-level normalization control. Negative control constructs were generated by introducing a premature stop codon upstream of the ribosomal slip (5’-termination), or downstream of a predicted -1 shift (3’-termination), a slip site mutant construct which disrupts the U_UUU_UUA slip site, and a positive control which contains an in-frame GFP cassette. Topological reporters were constructed by inserting a glycosylatable GFP (gGFP) cassette in-frame at the defined positions within the SINV structural polyprotein. Similarly, downstream of the test construct, an IRES-mKate was added as a normalization control.^19^

### Generation of CRISPR Knockout Cell Lines

CRISPR Knockouts lacking subunits of the multipass translocon were generated by the Genomics and Genome Editing Center at the Bindley Bioscience Institute in the context of a genetically modified HEK293T cell line containing two bio-orthogonal genomic Bxb1 recombination sites.^33^ CRISPR knockout cells were generated in a similar manner as previously described.^34^ Briefly, cells were transfected with SpCas9 enzymes that were pre-loaded with guide RNAs targeting each multipass subunit genes (Synthego, Redwood City, CA USA, see Table S2). Individual cells were then sorted into 96 well cell culture dishes then grown to confluency, harvested, and banked in liquid nitrogen for preservation. To validate the genomic edits, the genomic DNA was isolated from each clone prior to PCR amplification and Sanger sequencing of the target gene. Indels were then identified using the Synthego ICE analysis platform. Finally, the loss of the target protein was validated by western blot and whole-cell proteomics (Figs. 2 and S1).

### Whole-Cell Proteomics

Briefly, approximately one million cells from each knockout line were lysed by incubation in a Radio Immuno-Precipitation buffer (RIPA) containing SIGMAFAST protease inhibitor cocktail and Phosphatase Inhibitor Cocktail 2 (Sigma Aldrich, St. Louis. MO, U.S.A). Lysates were then clarified by centrifugation at 20,000 × g for 20 minutes. Protein concentrations of the supernatants were then determined using a BCA assay. Internal enolase standards were then added into 50 μg of protein from each sample to a final concentration of 0.5 μg/ μL prior to reduction of disulfide bonds through the addition of 5 mM dithiothreitol (DTT). Free cysteines were then alkylated by the addition of 15 mM iodoacetamide (IAA). Proteins were then digested overnight by Trypsin/LysC in an S-Trap− Mini Spin Column (ProtiFi, Fairport, NY, USA). Digested proteins were then desalted using a C18 column and resuspended in 5% acetonitrile containing 0.1% Formic Acid. Digested peptides were then analyzed using a timsTOF HT Mass Spectrometer (Bruker Corporation, Billerica, MA, USA) coupled to an inline nanoElute 2 liquid chromatography (LC) system. LC separation was carried out using the DIA-PASEF mode. To ensure precision across replicates, the system was calibrated using the Agilent ESI-L tuning mix and HeLa cell lysate standards. Digestion efficiency was validated based on the digestion of the internal enolase peptides using Skyline. Raw proteomic data were analyzed using spectronautV21 with a human spectral library. VSN normalization was applied, and the missing values were imputed using the Mindet function within the Analyst platform.

### Genetic Reporter Measurements

Reporter constructs were transiently expressed using Lipofectamine 3000 (Invitrogen, Waltham, MA, USA) in accordance with the manufacturer’s instructions. Cells were harvested 2 days post-transfection, washed twice with 1X PBS and analyzed using on a Attune NxT Flow Cytometer (ThermoFisher Scientific, Waltham, MA, USA). Cell fluorescence profiles were analyzed using FlowJoX software (Treestar Inc., Ashland, OR, USA). Fluorogenic control constructs were included in accordance with prevailing standards for bicistronic reporter constructs. Importantly, we previously ruled out artifacts that could arise from aberrant splicing of the reporter transcript in the context of HEK293T cells.^35^

### Frameshifting Efficiency Calculations

To calculate frameshifting on a representative sample of cells, they were restricted to include only cells with an IRES-mKate fluorescence within ± 2SD of the average median of mKate fluorescence of mKate+ cells. The mean fluorescence intensity of GFP and mKate of each construct was corrected for background by subtracting the fluorescence of the mKate-cell population. Frameshifting efficiency was calculated using,

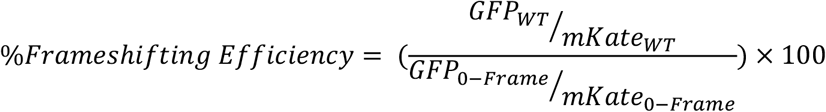

### Topology Reporter Calculations

To compare changes in frameshifting topology across MPT KO cell lines, cells were restricted to include only cells within ± 2SD of the average median of mKate fluorescence of mKate+ cells, per construct. The median fluorescence intensity for GFP and mKate was corrected by subtracting background signal. The corrected GFP/mKate ratio for each cell line was divided with the same construct in parental cell lines, to obtain relative GPF/mKate ratio.

### Preparation of Viral Stocks and Viral Titer Determination

To generate infectious Sindbis virus TE12-mCherry stocks, a cDNA clone encoding the viral genome was linearized by digestion with SacI (New England Biolabs, Ipswich, MA, USA). The linearized DNA served as a template for in vitro transcription to produce infectious viral RNA, which was transfected into BHK cells using Lipofectamine LTX according to the manufacturer’s instructions (Thermo Fisher Scientific, Waltham, MA, USA). To improve transfection efficiency and minimize serum interference, cells were transfected in Opti-MEM medium (Gibco, Grand Island, NY, USA). After 4 hours, the transfection medium was replaced with minimum essential medium (MEM) supplemented with 10% fetal bovine serum (FBS; Corning, Corning, NY, USA). Virus-containing supernatants were harvested 48 hours post-transfection, and cellular debris was removed by centrifugation.

Viral titers were determined by standard plaque assay using BHK cells. Briefly, serial dilutions of each viral stock were added to BHK cell monolayers at approximately 70–90% confluency in 6-or 12-well plates and incubated for 1 hour at room temperature with gentle rocking to facilitate viral adsorption. Following adsorption, cells were overlaid with a 1:1 mixture of 2% low-melting-point agarose and 2× MEM supplemented with 10% FBS. At 48 hours post-infection, cells were fixed with formaldehyde for 1 hour and stained with crystal violet to visualize plaques. Viral titers were calculated as plaque-forming units per milliliter (PFU/mL) based on the number of plaques observed and the corresponding dilution factor.

### Polyprotein Processing Measurements

Parental and knockout cell lines were infected with virus at an MOI of 1 and harvested at 48 hours post-infection (hpi). Cell pellets were resuspended directly in SDS-PAGE sample buffer and lysed. Viral proteins were separated by SDS-PAGE on 10% polyacrylamide gels and transferred to nitrocellulose membranes. Membranes were blocked with Intercept Blocking Buffer (LI-COR Biosciences, Lincoln, NE, USA) for 1 hour at room temperature and then incubated with rabbit polyclonal antibodies against SINV E1 and E2 proteins (Josman, LLC, Napa, CA, USA) at a dilution of 1:2,500. Primary antibody incubation was performed either overnight at 4°C or for 1 hour at room temperature. Following incubation, membranes were washed with PBS containing 0.1% Tween-20 (PBS-T) and incubated with IRDye 800CW goat anti-rabbit IgG secondary antibody (1:10,000; LI-COR Biosciences, Lincoln, NE, USA) for 1 hour at room temperature. After secondary antibody incubation, membranes were washed with PBS-T followed by a final rinse with Milli-Q water. Blots were imaged using an Odyssey CLx Near-Infrared Imaging System and analyzed with Image Studio software (LI-COR Biosciences, Lincoln, NE, USA). Relative cleavage efficiency is intensity of (E2/E1)/(p110+E2/E1).

### Viral Infection Measurements

Sindbis TE12-mCherry virus spread was monitored at each indicated MOI by tracking the increase in the mCherry signal across the surface areas over time using an IncuCyteS3 live-cell imaging system (Sartorius AG, Göttingen, Germany). Briefly, each cell line was seeded in a 48-well plate four hours before the start of infection. To account for variations in doubling time of the knockout cell lines, all cells were counted and seeded on the same day the experiment was performed. Viral inocula were then added to the wells and allowed to adsorb to the cellular monolayer for 1 hour at room temperature with gentle rocking. The plates were then placed in the IncuCyteS3 instrument for real-time monitoring of viral spread. Images were acquired every 3–6 hours over a 72-hour period at 10× magnification using both phase-contrast and red fluorescence channels. Four fields of view were imaged in each well of the 48-well plate at every time point. Image analysis was performed using Incucyte Base Analysis Software in Base Analyzer Mode. No segmentation adjustments were applied to the phase-contrast images, and the red fluorescence threshold was set between 20 and 30. To account for differences in cell confluency, the percentage of red fluorescent cells relative to total phase-detected cells was calculated and plotted.

## Supporting information

Table S1 Proteomic Analysis

## Acknowledgements

We thank Robert Keenan and Mukhopadhyay lab for helpful scientific input. We thank Wen-Hung Wang and the Genomics and Genome Editing Center at the Bindley Bioscience Institute for technical assistance with the generation of knockout cell lines. This work was supported by funding from the National Institute of General Medical Sciences (R35152086 to J.P.S.). The authors would also like to thank the National Institute of Health S10 instrumentation program for the funding support of the TIMS-TOF mass spectrometry (S10OD032364) and the Purdue Office of Research for the support of Acquisition of High-Field Asymmetric Waveform Ion Mobility Spectrometry. T.M. would like to acknowledge support from the Indiana University-Institute for Advanced Study collaborative research study award.

## Data Availability

All raw and processed experimental data are accessible through Mendeley data (doi: 10.17632/fhyxg5h6pk.2). Proteomic data can be found in the MassIVE repository (MSV000101782). Any other materials or information will be freely shared upon request.

## Notes

https://data.mendeley.com/research-data/?query=10.17632/fhyxg5h6pk.2

https://massive.ucsd.edu/ProteoSAFe/dataset.jsp?task=9c902819862a400fb12940a5d7040842

### Competing Interest Statement

The authors have declared no competing interest.

### Summary of Updates

There was an error in the order of the authors that were entered into bioRxiv. We have simply changed the order of the authors on the bioRxiv system so that it is consistent with the manuscript and reflects the author contributions. There are no material changes to the manuscript or supplement.

