## Supplemental Materials for "Translocon Remodeling Modulates Ribosomal Frameshifting and the Maturation of the Sindbis Virus Structural Polyprotein"

‡Authors contributed equally
*Corresponding authors: jschleba (at) purdue.edu, sumukhop (at) iu.edu

Contents:

-Figure S1
-Figure S2

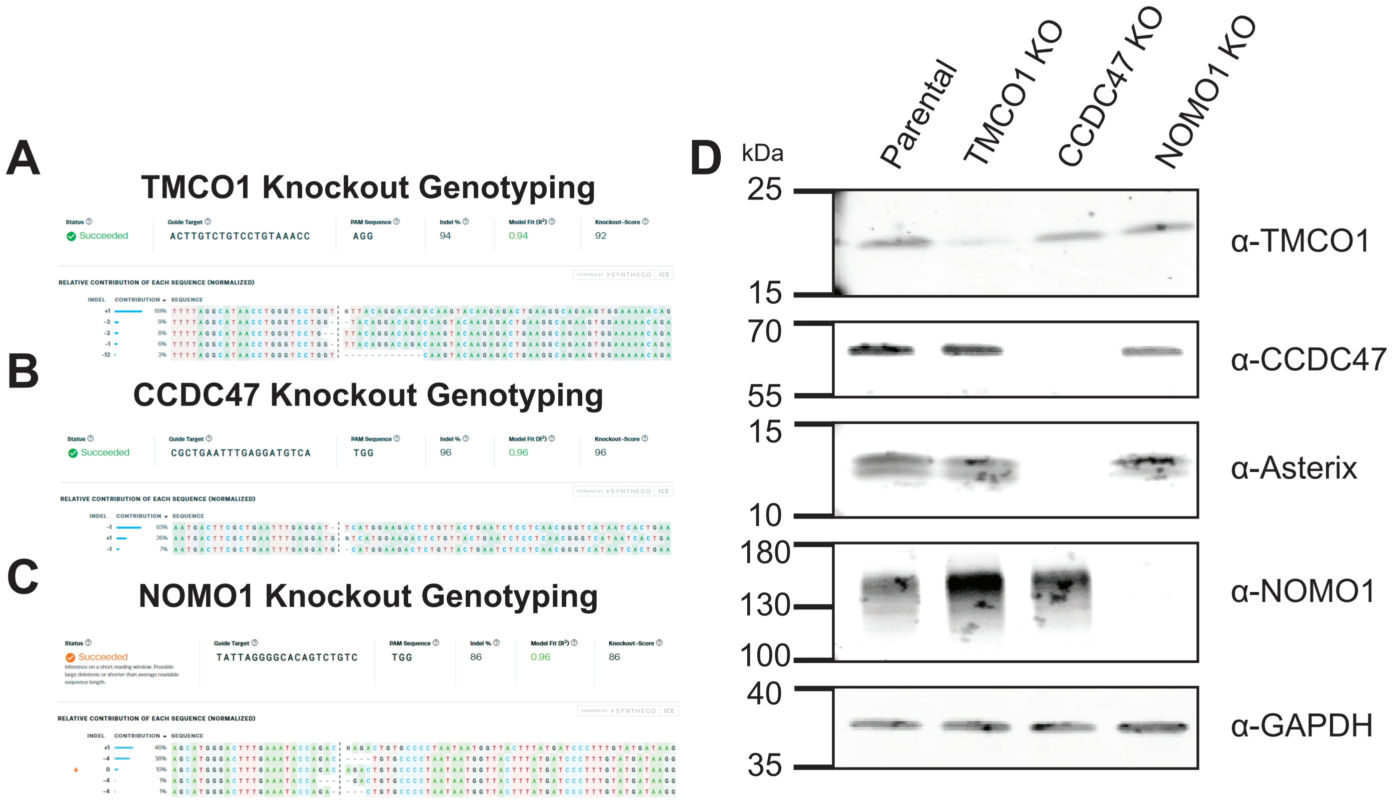


**Figure S1. Characterization of multipass translocon knockout cell lines.** Synthego ICE analyses of the genomic lesions within the A) *TMCO1*, B) *CCDC47*, and C) *NOMO1* genes of the corresponding CRISPR clones used in these studies are shown. The sequences of the sgRNA target and PAM, the inferred indel spectrum, the percentage of edited alleles (indel %), the model fit ($R^2$), and the knockout score for each clone are shown for reference. B) Loss of each target was confirmed using a western blot of parental and knockout lysates probed for TMCO1, CCDC47, Asterix, and NOMO1. A GAPDH loading control is shown for reference. The results are consistent with the trends observed using whole-cell proteomics (see Fig. S2 and Table S1).

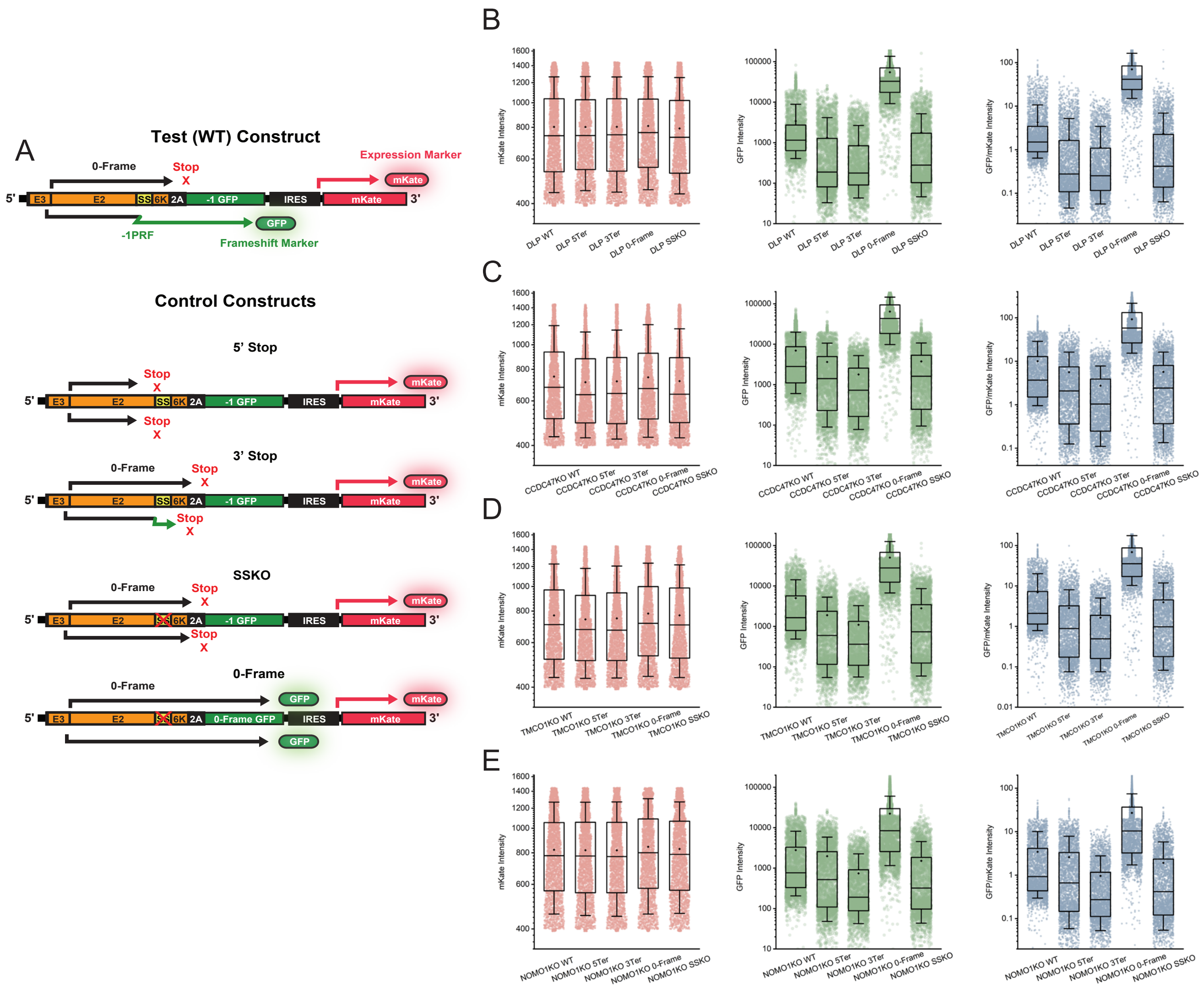


**Figure S2. Design of the fluorescent frameshifting reporter and controls.** A) Schematic of the bicistronic frameshifting reporter. The four control constructs are shown below: 5′Ter and 3′Ter carry premature and downstream stop codons, respectively; SSKO lacks the slippery sequence (slip-site knockout); and the 0-Frame construct encodes GFP in the 0 frame as a positive control. (B–E) Single-cell flow cytometry distributions of mKate intensity (left), GFP intensity (center), and the GFP/mKate ratio (right) for the WT, 5′Ter, 3′Ter, 0-Frame, and SSKO constructs in A) DLP Parental B), CCDC47 KO C), TMCO1 KO D), and NOMO1 KO E) cells. Each point is a single cell (n = [N] per condition); the box spans the interquartile range, the center line is the median, the dot is the mean, and whiskers indicate [5th–95th percentile / 1.5× IQR]. Vertical axes are shown in log scale.

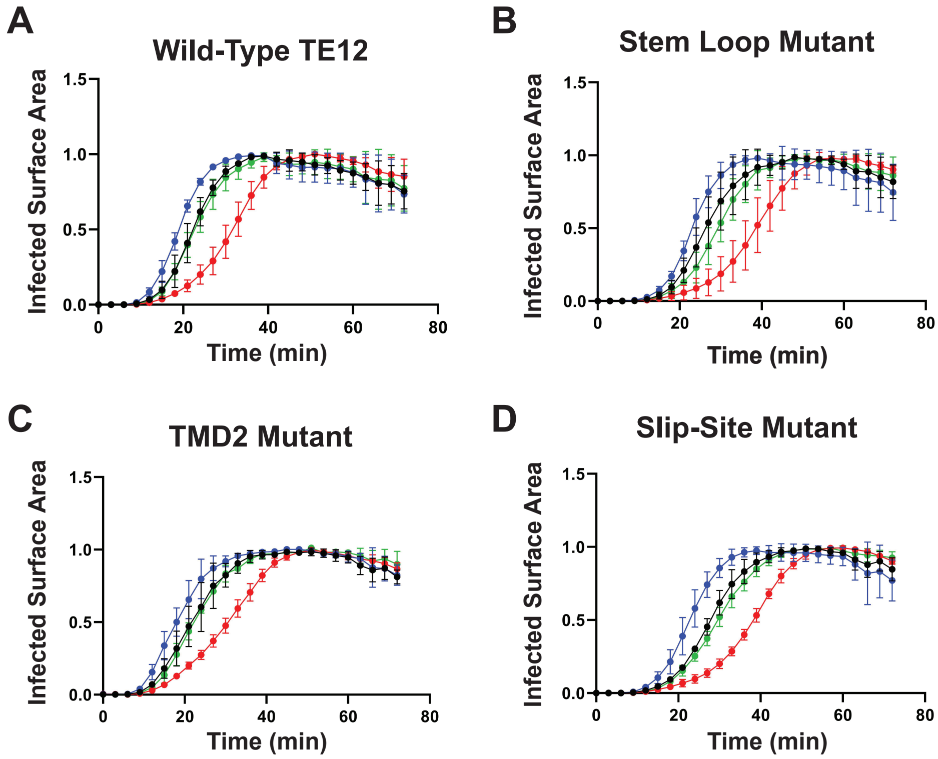


**Figure S3. Impact of MPT Depletion on the Fitness of SINV mutants with Impaired Frameshifting.** An incucyte was used to track the infection kinetics of several cell lines including TMCO1 knockout (blue), CCDC47 knockout (red), NOMO1 knockout (green), and parental HEK293T cells with A) WT TE-12-mCherry SINV and various other strains bearing mutations that disrupt -1PRF elements including B) the stem-loop (A2414G), TMD2 (V407E/ I408E), and the slip-site ($U_1$ $UUU_4$ $UUA_7$ to $G_1$ $UUC_4$ $CUA_7$). Cells were infected at MOI 0.2 and imaged every 3-6 hours for 72 hours by live-cell microscopy. Viral spread is expressed as virus area normalized to cell area (per image) with 4 images taken at 10x magnification per well in a 48-well plate. Growth kinetics in NOMO1 knockout cells were similar to the parent under these conditions, which may stem from differences in MOI. The fitness defect in CCDC47 knockout cells is consistent across mutated variants, suggesting it does not arise from changes in -1PRF efficiency.
